# RetS senses neurotransmitter norepinephrine and adrenergic agents to regulate *Pseudomonas aeruginosa* virulence

**DOI:** 10.1101/2024.11.24.625011

**Authors:** Yachung Zhou, Anming Ren, Chenchen Wang, Tian Zhou, Jie Deng, Yingfeng Huang, Ruiqin Cui, Wei Huang, Youjun Feng, Weidong Zheng, Haihua Liang, Liang Yang

## Abstract

Bacterial cells employ diverse mechanisms to perceive and respond to host-associated molecules, coordinate intracellular signal transduction, and engage in interkingdom communications. In this study, we firstly discovered that the adrenergic antagonist dronedarone (DDR) effectively suppresses quorum sensing (QS) and pathogenesis of the opportunistic pathogen *P. aeruginosa.* Further analysis revealed that DDR targets RetS, an bacterial surface receptor critical for suppressing the GacS/GacA two-component regulatory system (TCS). Interestingly, the adrenergic agonist norepinephrine (NE) was found to exert opposite effects against RetS, thereby reversing DDR-induced phenotypes in *P. aeruginosa*. Moreover, we identified the hinge region of RetS^DISMED2^ serves as a binding pocket of NE and DDR, which provides potential mechanisms of the antagonistic effect of NE and DDR. Our study demonstrated that NE promotes *P. aeruginosa* infections while DDR can serve as an anti-virulence compound against *P. aeruginosa* infections. Thus, RetS might serve as a bacterial adrenergic receptor which keeps the NE under constant surveillance.

**Significance:** Sensing and recognition of host-derived molecules, as well as therapeutic drugs, are crucial for the survival of pathogens during infection process. The adrenergic drugs are well known to affect virulence and pathogenesis of the opportunistic pathogen *Pseudomonas aeruginosa* (*P. aeruginosa*). In this work, we have provided in-depth characterization of a Norepinephrine (NE) receptor in *P. aeruginosa*, the RetS membrane protein. NE is the-adrenergic agonist known as a stress hormone, exerts its activity through interaction with mammalian adrenergic receptors (AR). It has been shown that increased NE concentration during early sepsis can stimulate the growth of various Gram-negative and Gram-positive bacteria, including *P. aeruginosa*. We revealed that both adrenergic agonist NE and another adrenergic antagonist dronedarone (DDR) are able to interfering the RetS-GacS/GacA signaling cascade of *P. aeruginosa* and execute opposite impact on the pathogenesis of *P. aeruginosa* during infections.

## INTRODUCTION

Sense and recognize host-derived molecules is a crucial strategy employed by bacterial pathogens to coordinate interactions between eukaryotic hosts, their associated symbionts as well as commensals (1). To cope with environmental changes and engage in a dynamic lifestyle, bacteria have evolved multiple levels of sensing systems, including cell-surface signaling systems, QS systems, nucleotide signaling networks (2–3). The TCS typically comprises a histidine protein kinase and a response regulator protein. This histidine protein kinase can be stimulated by specific environmental cues, leading to autophosphorylation at a histidine residue. Subsequently, the high-energy phosphoryl group is transferred to an aspartate residue in the response regulator protein, resulting in its activated confirmation for subsequent regulatory functions. Interestingly, diverse forms of TCS have been identified across different bacterial species. For instance, the RetS-GacS/GacA system has been extensively examined in *P. aeruginosa* for its critical role in controlling adaptation and pathogenesis (4–5). The surface sensor RetS has transmitter phosphatase activity against the receiver domain of GacS-P and takes phosphoryl groups from GacS-P. GacS phosphorylates the cognate response regulator GacA. GacA-P activates transcription of regulatory RNAs including RsmY and RsmZ, which sequester the translational regulator, RsmA, thereby regulating downstream genes and multiple phenotypes such as motility, QS, virulence and cyclic di-GMP signaling (5–9).

*P. aeruginosa* is a ubiquitous opportunistic pathogen responsible for a variety of diseases, including pneumonia, otitis media and urinary tract infections. It predominantly affects burn patients and individuals with compromised immune systems (10). Despite receiving adequate antibiotic treatment, persistent infections and high mortality rates are consistently observed in *P. aeruginosa* infection (10–11). Moreover, *P. aeruginosa* is recognized as a constituent of the ESKAPE group (*Enterococcus faecium*, *Staphylococcus aureus*, *Klebsiella pneumoniae*, *Acinetobacter baumannii*, *P. aeruginosa* and *Enterobacter sp*) owing to its virulence factors and remarkable capacity for antibiotic resistance. (12). Similar to other pathogens, *P. aeruginosa* employs diverse regulatory mechanisms, such as QS, cyclic di-GMP signaling and TCSs, to orchestrate its adaptation to the host environment and facilitate the production of virulence factors (1,13). Extensive research has been conducted to elucidate the operational mechanism of RetS-GacS/GacA networks (5,14,15), however, only a limited repertoire of host-derived molecules have been identified as cues capable of modulating these networks.

The interactions between eukaryotic hormones and pathogens, known as “microbial endocrinology” have been demonstrated to play significant roles in the context of infections (16, 17). For example, sex steroids such as testosterone and estriol have been documented to elicit membrane stress responses and *pqs* quorum sensing in *P. aeruginosa* (18). NE, a stress hormone and α-adrenergic agonist, exerts its effects through interaction with mammalian adrenergic receptors (AR). NE is commonly used for vasopressor therapy in septic shock cases. However, it has been demonstrated that elevated NE levels during early sepsis can promote the growth of various Gram-negative and Gram-positive bacteria, including *P. aeruginosa* (19–20). Stolk et al. reported that NE serves as an intermediary factor linking sepsis severity to immunosuppression and ultimately drives immunosuppression in sepsis (21). Additionally, NE has been shown to inhibit natural killer cell cytotoxicity (22) and play a role in bacterial resistance (23). Moreover, several studies have documented the regulatory effects of NE on various phenotypes of *P. aeruginosa*, including virulence, motility, and pyoverdine production. However, the underlying molecular mechanism behind this effect remains unknown (24–27). Revealing the direct impacts of NE on *P. aeruginosa* during infections is of paramount importance in unraveling the significance of microbial endocrinology.

In the present study, we initially discovered that DDR (28), a beta-adrenergic receptor antagonist, exerts diverse effects on various phenotypes of *P. aeruginosa*, including QS, virulence factor production, iron siderophore pyoverdine synthesis, motility, and intracellular survival. Subsequently, through molecular genetic analysis, we identified RetS as a surface sensor of DDR in *P. aeruginosa*. Interestingly, NE was also revealed as a ligand for RetS and can reverse or neutralize the effects of DDR on *P. aeruginosa*. Significantly, employing isothermal titration calorimetry (ITC) analysis, we have quantified the binding affinity between both DDR and NE with RetS, wherein NE exhibits a stronger binding affinity compared to DDR. Moreover, NE can establish a hydrogen bond network with residues from the hinge loop and its surrounding region of RetS, thereby further enhancing the stability of their conformation and might ultimately promote the monomeric form of RetS and block its interaction with GacS. Our data collectively demonstrate that RetS functions as a bacterial adrenergic receptor, regulating *P. aeruginosa* infection through the GacS/GacA two-component system. Furthermore, we provide an in-depth discussion on the underlying structural mechanism of RetS’s dual functionality.

## RESULTS

### DDR regulates QS and motility phenotypes in *P. aeruginosa*

QS is an abundant bacterial cell-to-cell communication mechanism that orchestrates interspecies and intraspecies interactions via diverse difusible chemical signals. It governs bacterial gene expression in a density-dependent manner, playing an indispensable role in the production of virulence factors (29). In our in house lab scale screening for QS inhibitors from FDA approved drugs (30), we discovered that DDR exhibits dose-dependent QS inhibition activity, effectively suppressing the expression of all three QS bioreporter genes (*lasB*-*gfp*, *pqsA*-*gfp* and *rhlA*-*gfp*) in *P. aeruginosa* (Fig. 1*A*), indicating that DDR targets a general regulatory mechanism governing QS. In consistent with report fusion results, DDR was observed to suppress the synthesis of QS-dependent virulence factors, including elastase, pyocyanin, rhamnolipids (Fig. 1*B*), and partially QS-dependent virulence factor pyoverdine (Fig. S1). Furthermore, DDR significantly enhanced the motility (Fig. S2) of *P. aeruginosa* without any discernible impact on growth enhancement (Fig. S3).

**Fig. 1.**
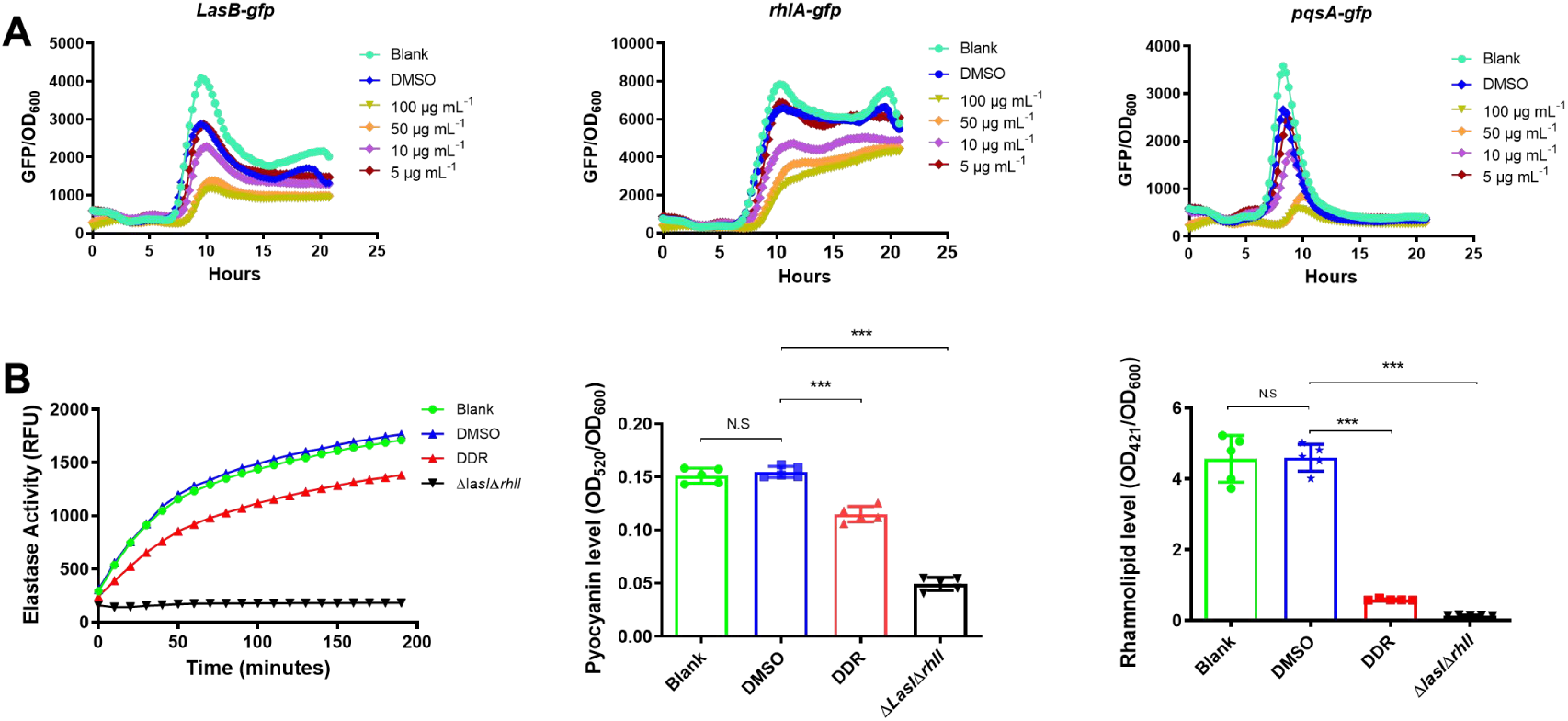
Effects of DDR on QS and the production of virulence factors in *P. aeruginosa*. (***A***) Dose-dependent inhibitory curves of DDR on QS monitor strains including PAO1-*lasB*-*gfp*, PAO1-*PqsA*-*gfp* and PAO1-*rhlA*-*gfp*. (***B***) The impact of DDR (50 μg mL^-1^) on QS regulated virulence factors including elastase, pyocyanin, and rhamnolipid, QS defective PAO1-Δ*lasI*Δ*rhlI* mutant was used as a negative control. One way ANOVA was employed for significance evaluation, error bars means ± SDs, NS means no-significant, *** denotes p < 0.001.

**Fig. S1.**
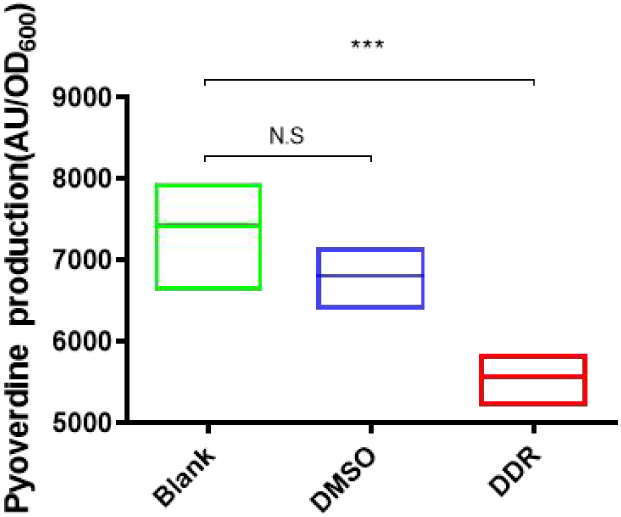
Effect of DDR on *P. aeruginosa’s* pyoverdine production. Infulence of DDR treatment on the production of pyoverdine. One way ANOVA was employed for significance evaluation, error bars means ± SDs, NS indicates no-significant, *** denotes p < 0.001.

**Fig. S2.**
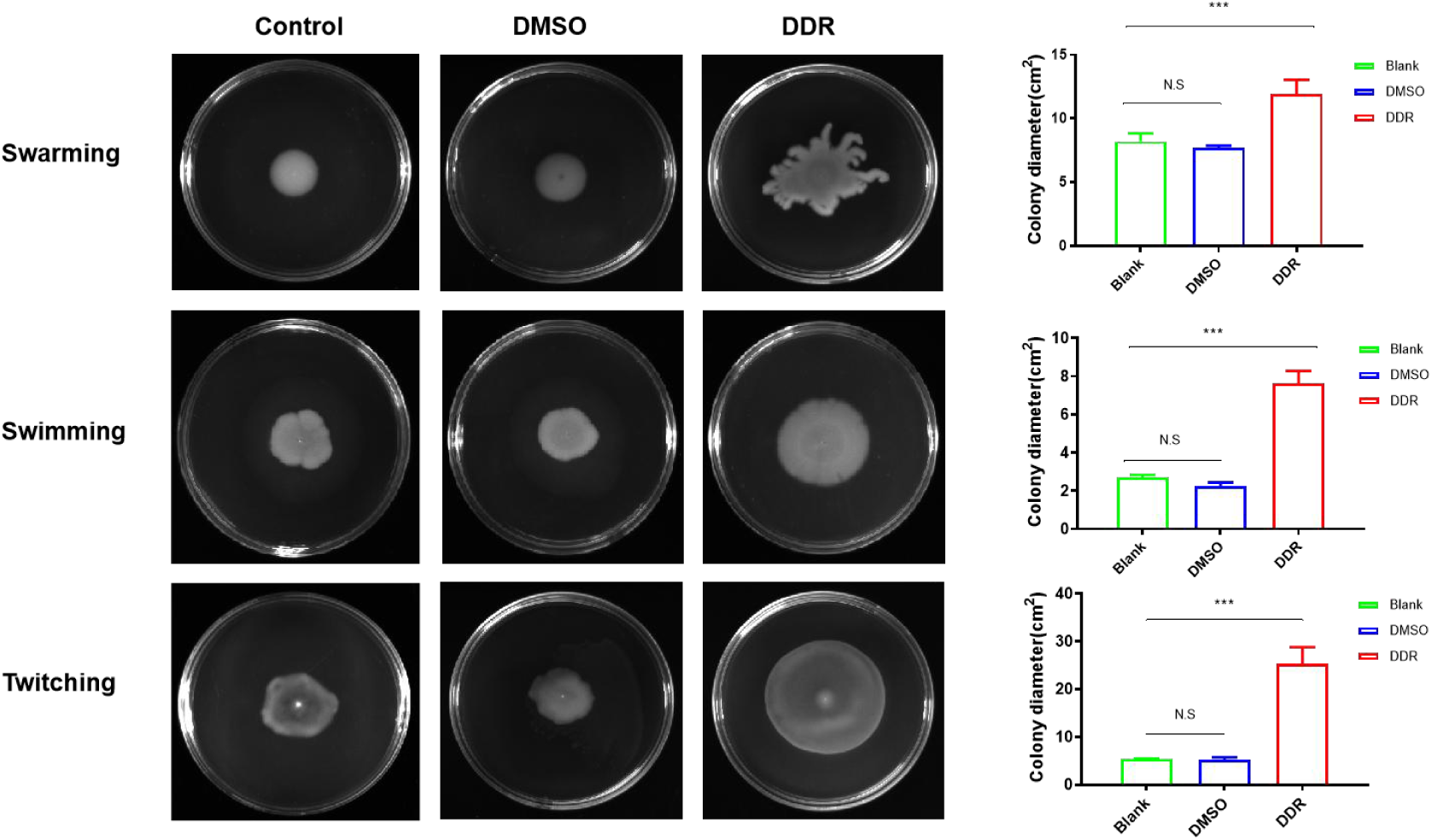
Impact of DDR on *P. aeruginosa’s* motility. Effect of DDR (50 μg mL^-1^) on *P. aeruginosa’s* motility after incubation with 50 μg mL^-1^ DDR, DMSO or blank control in motility assay plates including swarming, swimming and twitching.

**Fig. S3.**
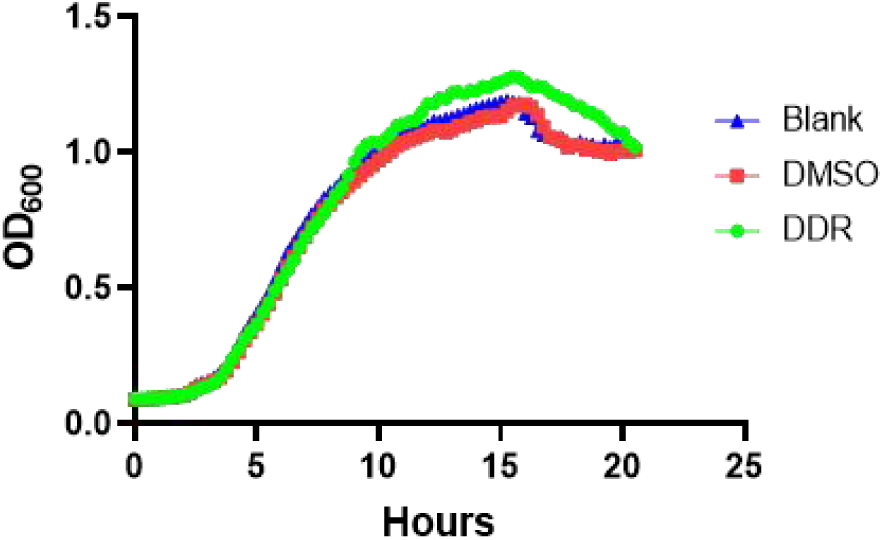
Effect of DDR (50 μg mL^-1^) on *P. aeruginosa’s* growth.

### DDR reduces pathogenesis of *P. aeruginosa* during infections

Given DDR’s potent ability to suppress the production of diverse virulence factors, we conducted further investigations into the impact of DDR on *P. aeruginosa* infections. As illustrated in Figure 2, RAW264.7 macrophages infected with *P. aeruginosa* at various multiplicity of infection (MOI) exhibited attenuated cytotoxicity (Fig. 2*A*) and reduced intracellular survival rates upon DDR treatment compared to the control group, although statistical significance was only observed for MOI ranging from 10 to 20 in the intracellular survival assay (Fig. 2*B*). Consistent results were obtained from the *Galleria mellonella* infection model (Fig. 2*C*). To gain deeper insights into the role of DDR in *P. aeruginosa* infection, a murine lung infection model was established, revealing that non-infected controls displayed intact single-layer cells surrounding alveoli (Fig. 2*D*). In contrast, in the infection group, DDR treatment significantly attenuated *P. aeruginosa* PAO1 pathogenesis compared to the blank control group, evident thickening of epithelial cell layers, massive recruitment of immune cells and destruction of alveolar septa were observed along with persistent leukocyte infiltration and severe edema (Fig. 2*D*). These data strongly suggests that DDR effectively inhibits pathogenesis and attenuates infection caused by *P. aeruginosa*.

**Fig. 2.**
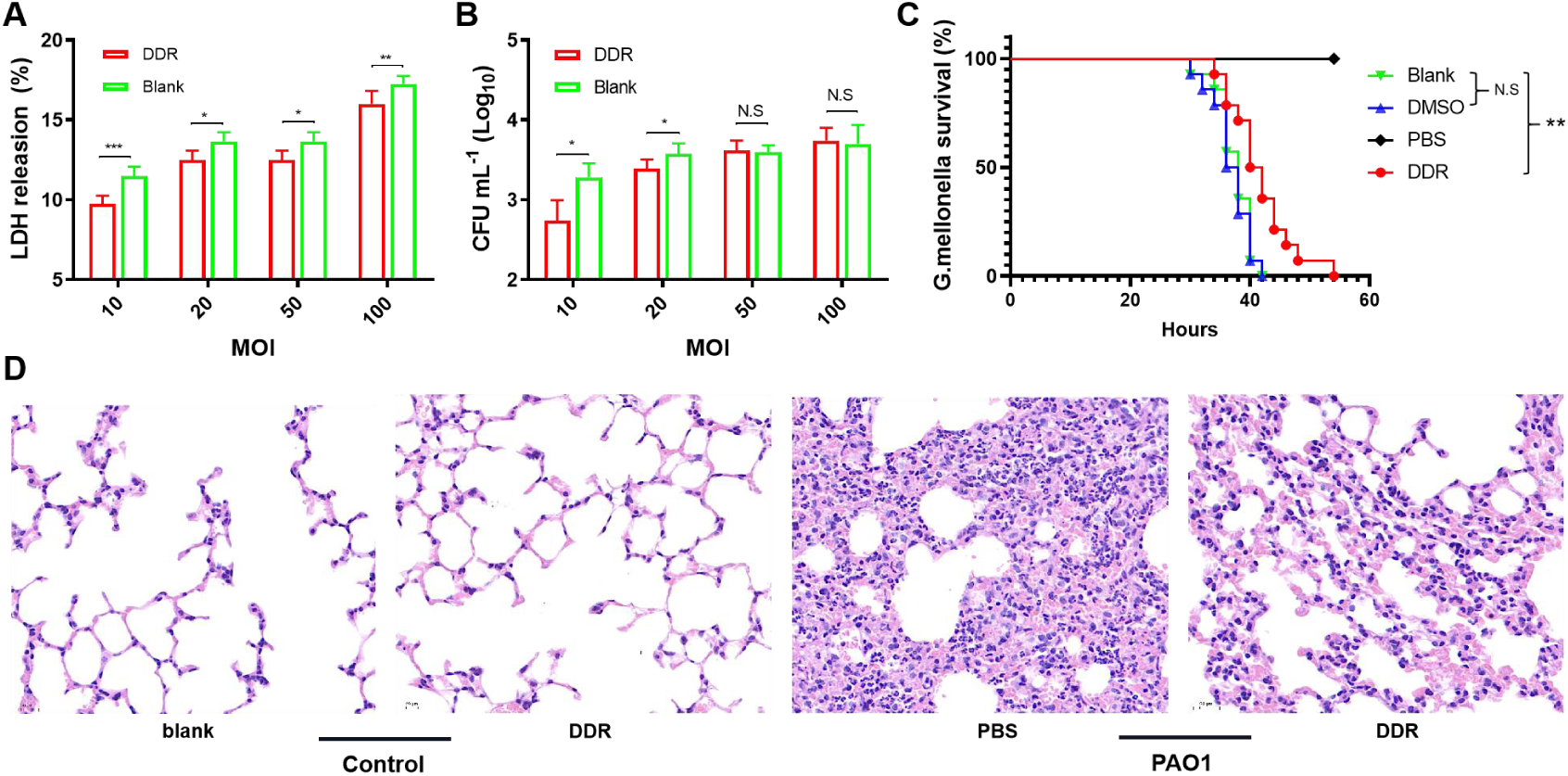
Impact of DDR on *P. aeruginosa* infection. (***A*-*B***) Virulence and intracellular viability of PAO1 cultured with 50 μg mL^-1^ DDR or not. Error bars represent means ± SDs. One way ANOVA was employed to evaluating significance (n=4), NS means no significance, * = p < 0.05, *** = p < 0.001. (***C***) The effect of DDR treatment on the survive rate of *Galleria mellonella* after *P. aeruginosa* infection. ** = p < 0.01, Kaplan-Meier was employed to conduct the statistic analysis (n=10). (***D***) Lung HE-stained sections of DDR pretreated mice that infected with *P. aeruginosa* or not.

### DDR modulates *P. aeruginosa* virulence phenotypes through the RetS-GacS/GacA network

Given the observed phenotypic changes in *P. aeruginosa* induced by DDR treatment, which are known to be regulated by the GacS/GacA TCS, we hypothesized that DDR targets this TCS (5, 31–34). To test our hypothesis, we employed qRT-PCR assay and several bioreporter strains of the RetS-GacS/GacA networks, including PAO1-*rsmY*-*gfp*, PAO1-*rsmZ*-*gfp* and PAO1-*cdrA*-*gfp* [*cdrA* expression is routinely used as an indicator of intracellular c-di-GMP levels (35)], to investigate the impact of DDR on this TCS. As expected, DDR treatment upregulated *retS* expression while downregulating *gacA* expression in *P. aeruginosa*, resulting in increased *rsmA* expression (Fig. 3*A*) and activation of downstream transduction pathways, e.g. *rsmY* and *rsmZ* (Fig 3*B*-*D*). Importantly, DDR treatment failed to affect *P. aeruginosa* motility (Fig 3*E*) in the Δ*retS* mutant, and the QS phenotypes (Fig. 1*A*) were rescued in this mutant strain (Fig. 3*F*). Collectively, these data indicate that DDR regulates *P. aeruginosa* phenotypes through modulation of the RetS-GacS/GacA pathway, strongly suggesting that RetS serves as a putative target of DDR.

**Fig. 3.**
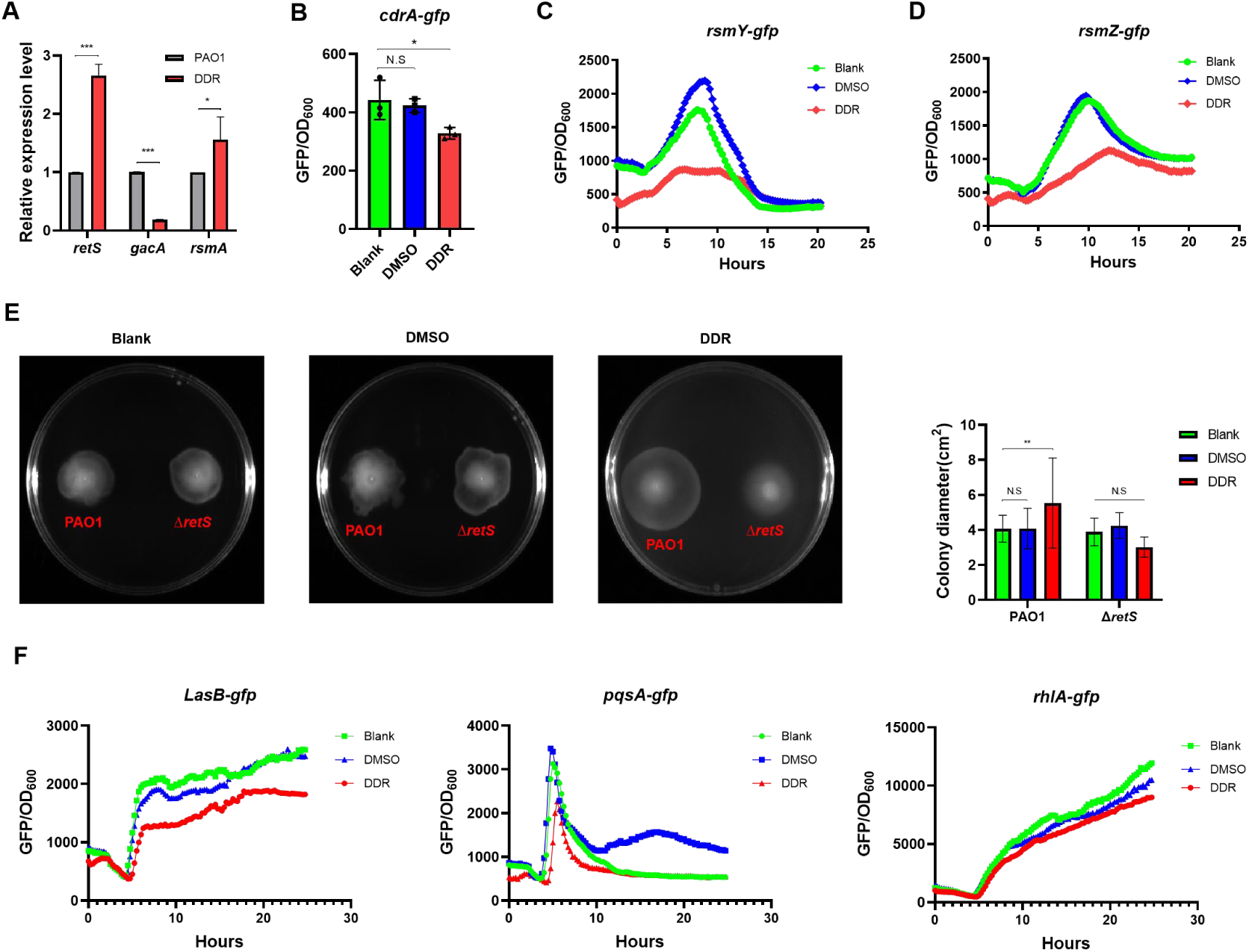
Impact of DDR on the *P. aeruginosa* RetS-GacS/GacA network. (***A***) RT-qPCR of RetS-GacS/GacA network-related genes after 50 μg mL^-1^ DDR treatment. Significance was evaluated using One way ANOVA (n=3), error bars represent means ± SDs. NS indicates no significance. * = p < 0.05. *** = p < 0.001. (***B*-*D***) Effect of DDR treatment (50 μg mL^-1^) on the reporter fusions of RetS-GacS/GacA network, including PAO1-*rsmY*-*gfp*, PAO1-*rsmZ*-*gfp* and PAO1-*cdrA*-*gfp*. (***E***) Swimming motility of wild type of PAO1 and Δ*retS* mutant in plates with DDR added or not. Statistical analysis was listed in the right side. ANOVA was used to evaluate significance (n=3), error bars indicate means ± SDs. NS means no significance. * *= p < 0.01. (***F***) Effect of DDR treatment (50 μg mL^-1^) on the expression of reporter fusions in the Δ*retS* mutant.

### DDR and NE target RetS sensor in *P. aeruginosa*

Clinically, DDR functions as an antagonist of beta-adrenergic receptors, while NE acts as an agonist of adrenergic receptors, thereby exhibiting reciprocal antagonism. As demonstrated above, DDR effectively modulates the gene expression of the RetS-GacS/GacA regulatory network. Given the fact that deletion of *retS* abrogated the effects of DDR on *P. aeruginosa*-related phenotypes, we postulated that RetS may represent a potential target for NE and DDR. To evaluate the binding of NE and DDR with RetS, we carried out an isothermal titration calorimetry (ITC) assay using the recombinant RetS fusion protein with both compounds. The signaling domain of hybrid sensor RetS consists of 7-transmembrane region (7TMR, corresponding to RetS189-389) and a periplasmic sensor domain (diverse intracellular signaling module extracellular 2, DISMED2, corresponding to RetS48-171) (36–39). Clearly, Figure 4*A* showed that there are direct contacts between RetSDISMED2 and NE (∼1μM) or DDR (∼3.6μM), highlighting a higher affinity to RetSDISMED2 of NE than that of DDR.

**Fig. 4.**
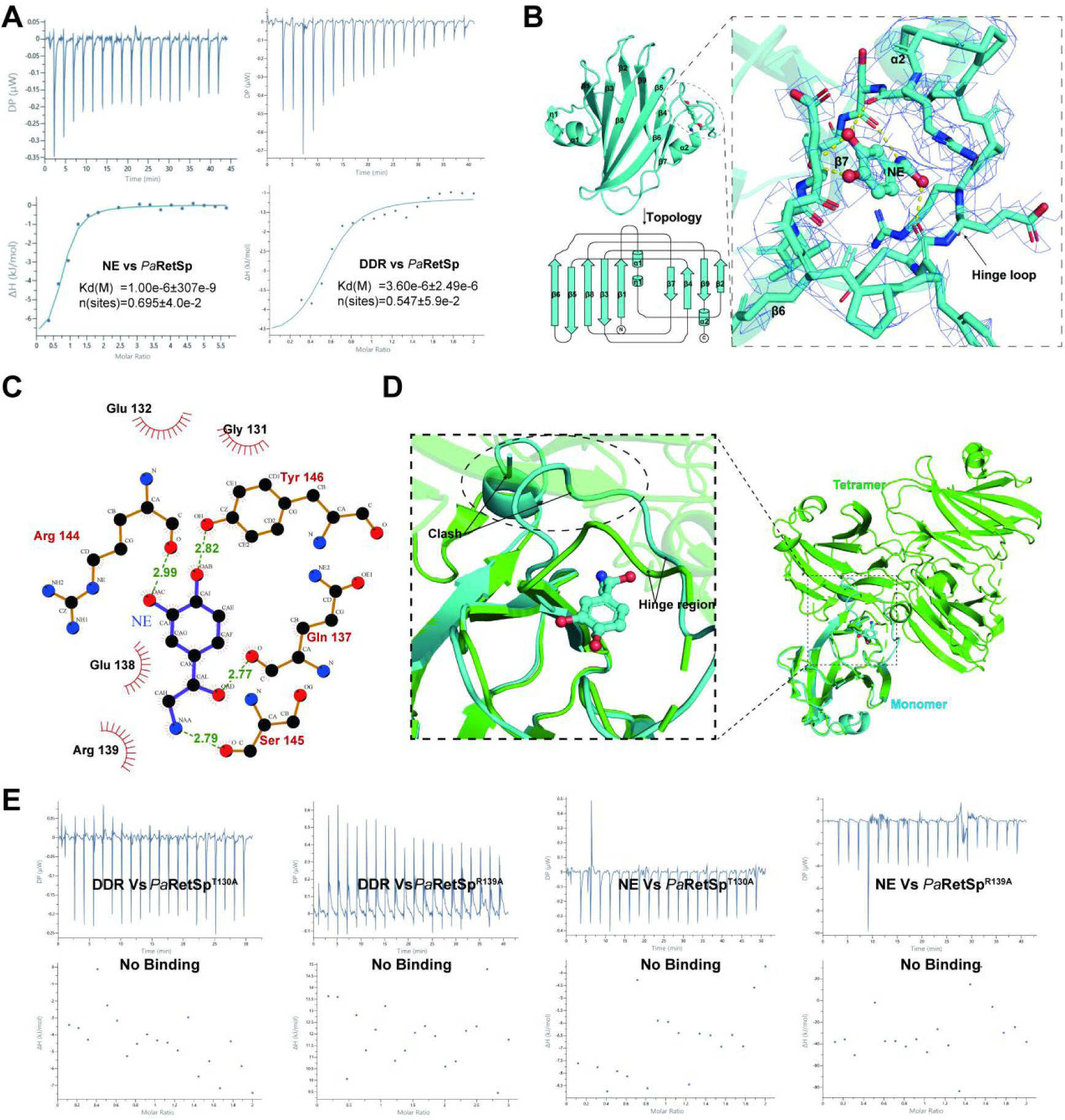
Ligand binding to the hinge region could regulate the conformation of PaRetSp hinge region. (***A***) ITC assays for evaluating the direct interactions between NE and DDR ligands and PaRetSp. (***B***) Cartoon diagram of PaRetSp (upper left) with the ligand pocket surrounded by an elliptical black dashed line and enlarged on the right panel. The electronic density map of hinge region and NE were represented with blue mesh contoured at 0.8σ. And the topology diagram of PaRetSp (bottom left) with each secondary structural element indicated in black. (***C***) 2-dimensional interaction diagram of NE bound to PaRetSp, which is rendered using Ligplot+ v2.2. **(*D***) Cartoon diagram of the superposition of monomeric PaRetSp with homo-tetramer PaRetSp highlighting the severe steric clashes between compact monomeric form and extended form observed in domain-swapped homo-tetramer form. **(*E***) ITC assays showing mutations of residues located in the hinge loop of *Pa*RetSp will lose the binding ability to ligands.

**Fig. S4.**
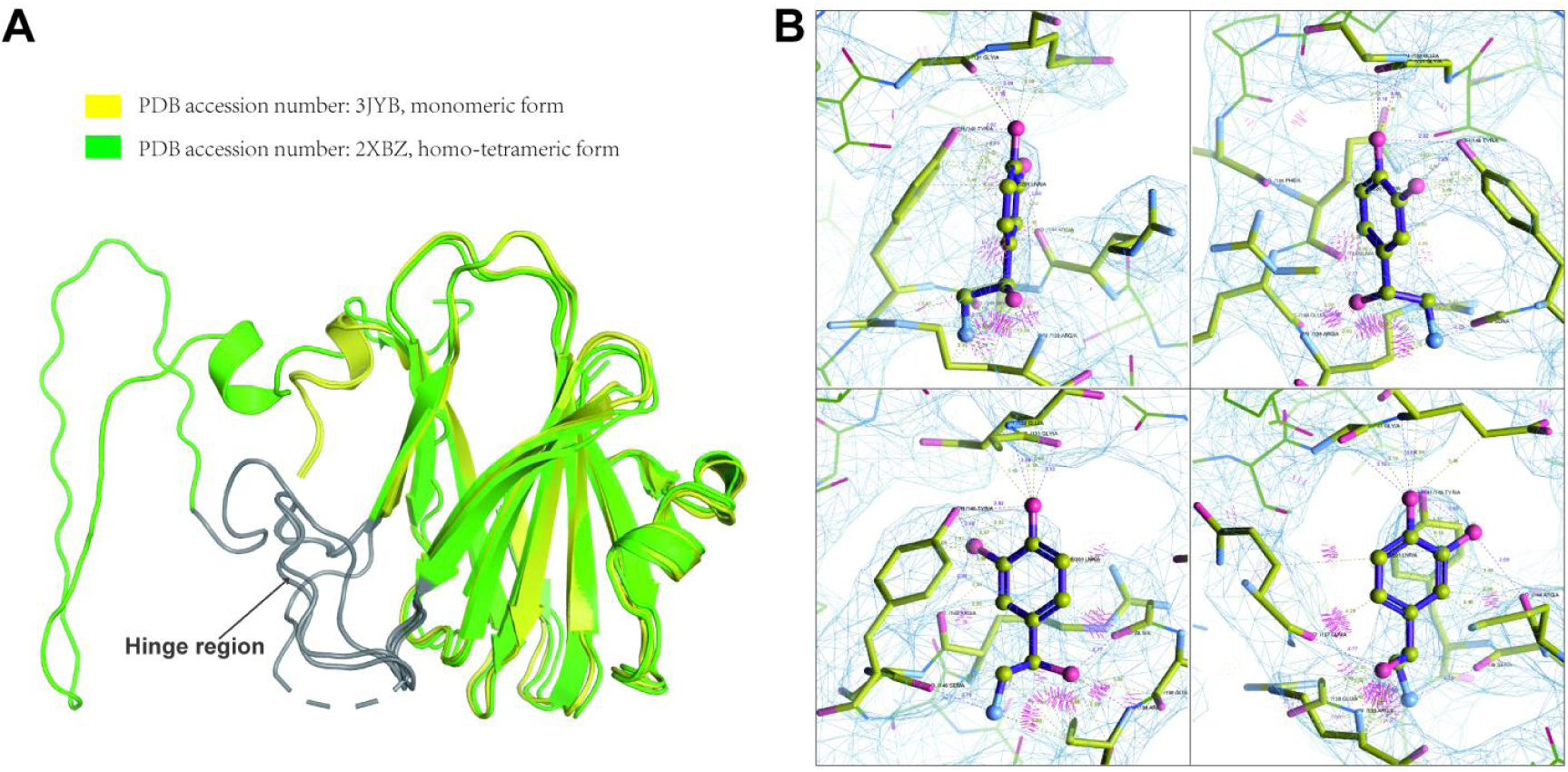
Ligand binding to the hinge region could regulate the conformation of *Pa*RetSp hinge region. (***A***) The cartoon diagram of the superposition of monomers extracted from crystal structures reported before with hinge regions colored in gray. And yellow ones were extracted from crystal structure of which PDB accession number is 3JYB; green ones were extracted from 2XBZ. (***B***) Several views of ligand NE with the electronic density map represented with blue mesh contoured at 0.8σ.

To gain a deeper insight into the structural basis of ligand binding details to RetSDISMED2 (referred as PaRetSp hereafter) and verify whether the domain-swapped form of which has any physiological relevance to its ligand binding activity, we crystallized PaRetSp in complex with NE solving the 3.5 Å resolution single crystal structure of PaRetSp. However, due to the low solubility of DDR and difficulty in crystallizing, we failed to solve the structure of PaRetSp-DDR. Previous study showed that there are two macromolecules in the asymmetric unit (AU) of its corresponding apo crystal forms, including the PaRetSp monomer (PDB accession code: 3JYB) with the symmetry of P212121 and its homo-tetramer (PDB accession code:2XBZ) with the symmetry of I4132 (40). In contrast, only one macromolecule could be observed in the AU of PaRetSp single crystal whose symmetry should be attributed to the crystallographic space group P43212. Moreover, as Figure S4A demonstrated, the hinge region presented intrinsic flexibility and variable conformation in the monomeric form extracted from above-mentioned two crystal structures, as reflected by higher B-factor or the lack of interpretable density over the hinge region in some certain monomer. And the hinge region plays a key role in the PaRetSp oligomerization, especially the stretch of residues Pro140-Leu141-Pro142-Ser143 from hinge loop, which contributes to the transition from the β turn and linear conformational forms of this hinge region (40). Even though the low resolution, all 124 amino acid residues (aa) of PaRetSp domain that adopts a classic jelly-roll fold, including hinge loop region (RetS130-145), could be well modeled into the corresponding well-defined electron density map. The discrepancies mentioned above hinted the entry of NE to the ligand-binding pocket located in PaRetSp. As expected, the small molecule NE could be fitted into an extra empty blob in the vicinity of hinge loop (Fig. 4*B* and Fig. S4*B*). NE was anchored to the pocket formed by the hinge region and strands β6 and β7. And Figure 4C shows that NE formed a hydrogen bond network with residues Q137, S145 and R144 located in the hinge loop together with R144 from strands β7. It could be inferred that due to the binding of NE, the PaRetSp hinge loop, whose flexibility is required by oligomerization (Fig. 4*D*), could be further stabilized. In addition, mutants (T130A and R138A) of PaRetSp suggested that DDR may occupy the same pocket (Fig. 4*E*), further participating in the regulation of the conformation of hinge region. However, T130A and R138A mutants of PaRetSp will also abolish the interaction between NE and PaRetSp (Fig. 4*E*). Taken together, NE could competitively bind to PaRetSp with DDR, further stabilizing the conformation of hinge loop referring to the domain-swapped form of PaRetSp observed in tetramer. Our data collectively demonstrated that DDR and NE target the RetS sensor of *P. aeruginosa*.

### NE could neutralize or reverse DDR regulated phenotypes in *P. aeruginosa*

Considering the direct binding of DDR and NE with RetS, it is evident that NE exhibits a larger impact on RetS. Consequently, we infer that NE may exert an antagonistic effect on DDR during *P. aeruginosa* infection. As expected, NE significantly reversed the stimulatory effect of DDR on swimming motility in wild-type *P. aeruginosa* but not in the Δ*retS* mutant (Fig. 5*A*). Moreover, NE effectively counteracted the inhibitory effects of DDR on virulence to a level comparable to the blank control in both *Galleria mellonella* larvae infection model (Fig. 5*B*) and murine lung infection model (Fig. 5*B*), while this phenomenon was absent in the Δ*retS* mutant group (Fig. 5*C*). Additionally, NE actively promoted pathogenesis of wild-type *P. aeruginosa* PAO1 but had no impact on the Δ*retS* mutant (Fig. 5*C*). We have noted that unlike in the *Galleria mellonella* infection model, Δ*retS* mutant was unable to cause severe pathogenesis in the murine lung infection model compared to PAO1, perhaps due to the repression of T3SS mechanism of *P. aeruginosa* of *retS* deletion (41). These findings are consistent with a previous study demonstrating that NE significantly enhances *P. aeruginosa* virulence (24). Besides, these data align well with the contrasting physiological functions of NE and DDR. Importantly, our findings strongly indicate that both NE and DDR target the RetS-GacS/GacA networks, thereby emphasizing NE’s role in promoting *P. aeruginosa* infections while highlighting DDR as a potential anti-virulence agent against such infections.

**Fig. 5.**
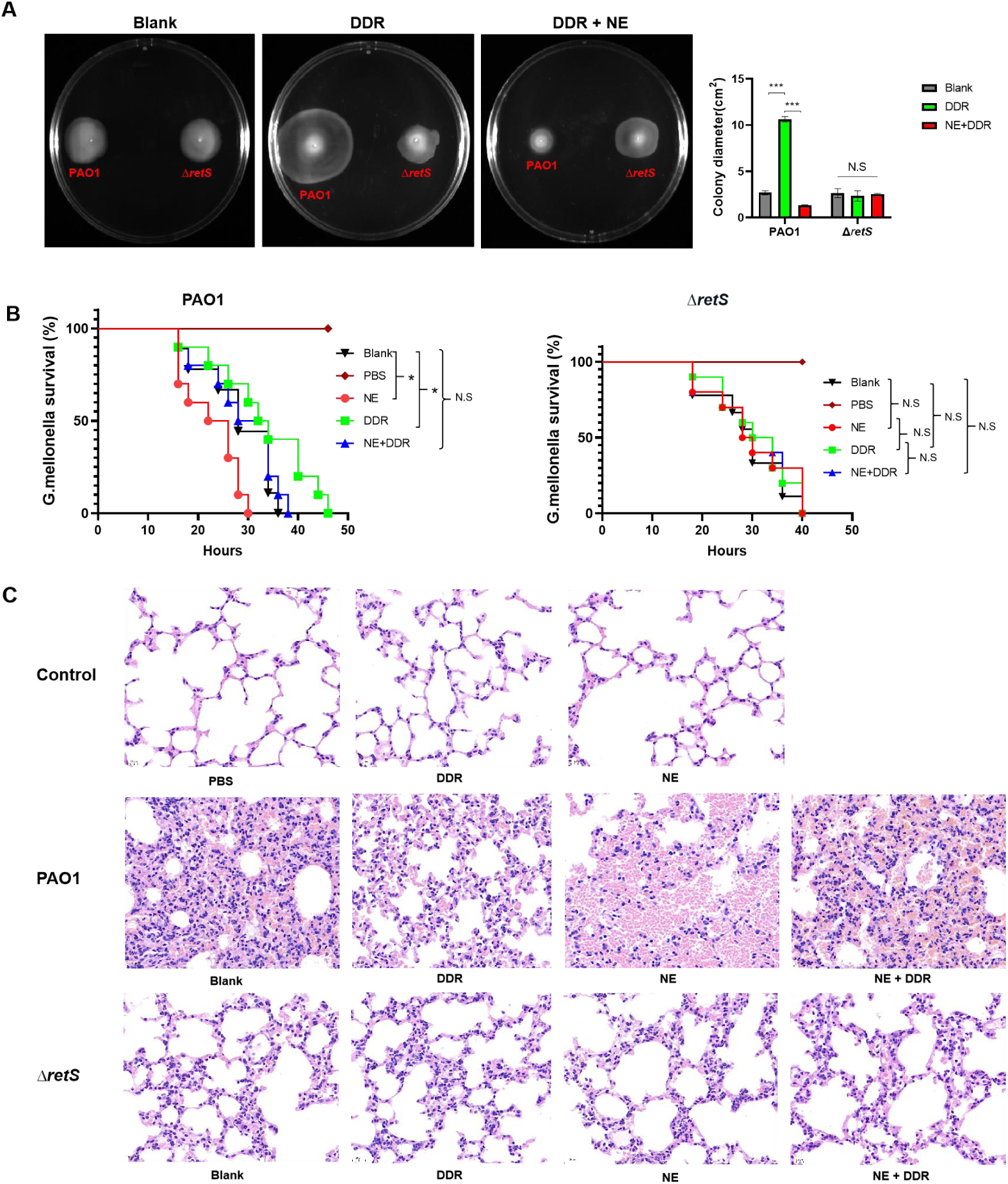
Antagonistic effect of NE on DDR-regulated phenotypes in *P. aeruginosa.* (***A***) The impact of NE and DDR on the swimming motility of both wild type PAO1 and Δ*retS* mutant strains was assessed by incubating them in plates containing DDR (50 μg mL^-1^) + NE (150 μg mL^-1^) or DDR alone (50 μg mL^-1^). A blank plate was used as a control. Statistical analysis using one-way ANOVA with a sample size of n=3 was performed to determine significance. NS means no significance. *** = p < 0.001. (***B***) The effect of NE and DDR on the survival rate of *Galleria mellonella* after PAO1 and Δ*retS* mutant infection, Kaplan-Meier was used to conduct the statistic analysis (n=10). (***C***) Representative lung histological images of mice that pretreated with NE or DDR followed by challenged with *P. aeruginosa* or Δ*retS* mutant.

## DISCUSSION

Norepinephrine (NE) belongs to the catecholamine amine hormones, which function as sympathetic neuroendocrine mediators in the “fight or flight” response to acute stress (42). The compound DDR exerts a potent inhibitory effect on multiple ion currents, including potassium current, peak sodium current, and L-type calcium current. Additionally, it demonstrates antiadrenergic properties (28). Previous studies have reported that physiological concentrations of NE in the gastrointestinal tract can reach as high as 50 mM (17). Furthermore, the gastrointestinal microbial system has evolved specific detection systems to sense host cues such as catecholamines and modulate their growth and virulence accordingly (43). It is well established that stress-induced molecules can significantly impact bacterial physiology (19). Despite several investigations on the effect of adrenergic agonists and antagonists on *P. aeruginosa*, an opportunistic pathogen commonly associated with hospital-acquired infections, the underlying mechanism remains elusive (24,25,27,44).

Given the extensive research on stress hormones, including NE, and their impact on diverse bacterial phenotypes (45–47), our study aims to investigate the molecular targets of adrenergic agents for their effects on various phenotypes of *P. aeruginosa*. Hegde et al previously reported that NE enhances virulence and motility in the PA14 strain of *P. aeruginosa*, potentially through modulation of the *las* QS (24). In this study, we not only demonstrated that DDR inhibits the *las* QS but also exerts a suppressive effect on two other QS systems: *pqs* QS and *rhl* QS systems. However, another study revealed contrasting findings where NE significantly suppressed pyoverdine production and expression of an important virulence factor (exotoxin A15 coded by toxA) instead of enhancing overall virulence production observed in other bacteria (27). Furthermore, Stolk et al emphasized NE as an intermediary factor that links sepsis severity to immunosuppression, ultimately driving immunosuppression in sepsis (21). Previous studies have demonstrated that catecholamines can enhance motility and colonization in *P. aeruginosa* PA14 (24). In contrast, our study revealed that DDR significantly enhanced three types of motility in *P. aeruginosa* PAO1. These contrasting effects of catecholamines and adrenergic receptor antagonists on bacterial virulence may be attributed to the modulation of diverse receptors involved in different regulatory networks within bacteria, while RetS may also respond to other host-derived molecules. Moreover, since NE has been reported to inhibit natural killer cell cytotoxicity (22), whereas destruction of noradrenergic nerve endings increased bacterial resistance in mice (23), it is plausible that the host immune response plays a role in the observed phenotypic differences among bacteria. Therefore, further investigations are warranted to elucidate the specific receptors and downstream signaling pathways involved.

As a fundamental principle, bacteria rely on presence sensors to detect signal molecules in their surrounding environment (48). Previous studies have demonstrated the existence of an adrenergic sensor in *S.enterica serovar Typhimurium* as well (49–50). Moreover, the TCS of QseBC and QseEF have been implicated in the response of *E. coli O157:H7* to NE (46,51). *P. aeruginosa* possesses multiple regulatory TCSs that contribute to its remarkable adaptability to diverse environmental conditions (52). In this investigation, we identified RetS as an adrenergic sensor that interacts with the GacS/GacA TCS and modulates phenotypes induced by NE and DDR through direct binding to RetS in *P. aeruginosa* PAO1. According to Florence et al., PaRetSp can exist as a dimer and a low-abundance tetramer in solution, along with the monomeric form. The dimeric PaRetSp predominates in solution, particularly at higher concentrations (7 mg mL^-1^). Furthermore, at lower concentrations (3.4-4.9 mg mL^-1^), the tetrameric form of PaRetSp, along with the monomeric and dimeric forms, can be observed (40). However, only the monomeric PaRetSp is capable of forming a heterodimer with GacS, thereby inhibiting GacS autophosphorylation and further interfering with related phynotypes (40,53,54). On the contrary, PaRetSp oligomerization will actually make an antagonistic effect on the related phynotypes (40). Regarding domain-swapped proteins, they typically exhibit a swapped domain either at the N- or C-terminus; however, only a limited number of cases have been reported where a portion of the macromolecule is observed to be swapped, as seen in PaRetSp (55). In the case of PaRetSp homo-tetramer, the domain-swapped dimer connects with two additional compact monomers to form a tightly-packed tetramer, which may hold significant biological relevance for RetS regulation. Furthermore, an extended domain-swapped dimer and a compact monomer are formed due to flexibility in the hinge loop region (40). Specifically, the hinge loop of PaRetSp plays a role in transitioning from its compact to extended form. The higher stability of Pro135 to Arg144 located within this hinge region is primarily compensated by hydrogen bond networks and van der Waals contacts within the oligomer (40). Our study provides novel insights into how catecholamine agonists (NE) and antagonists (DDR) modulate various phenotypes of *P. aeruginosa* by revealing RetS as a bacterial adrenergic receptor (Fig. 6).

**Fig. 6.**
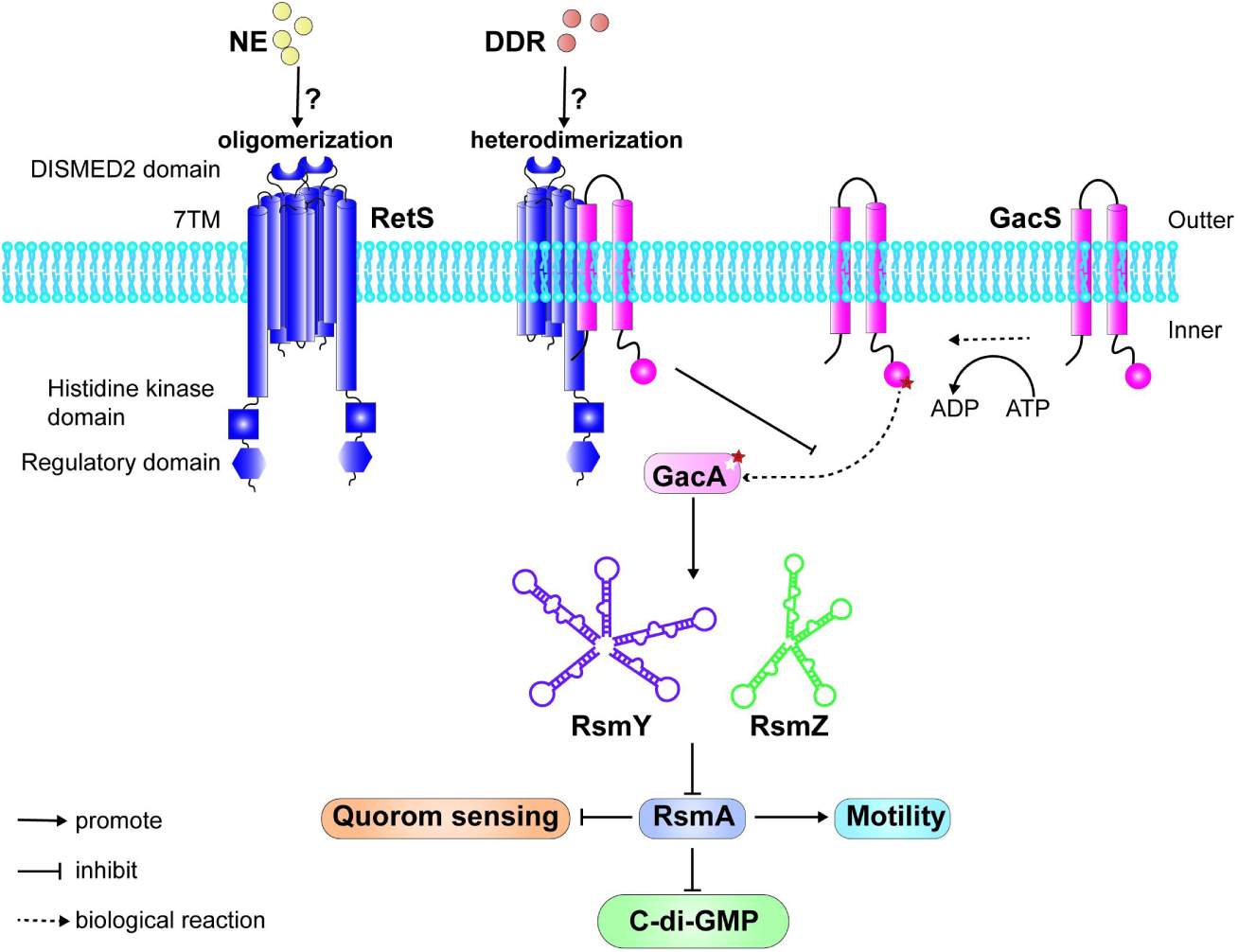
Diagram of NE and DDR regulate RetS-GacS/GacA system by directly binding with RetS sensor. NE and DDR modulate diverse virulence phenotypes of *P. aeruginosa* through the RetS-GacS/GacA system.

As discussed above, catecholamines and related adrenergic antagonists such as Timolol (TIM) and phentolamine exhibit diverse roles in various bacteria (24,25,44,45,60), however, the precise mechanisms underlying these phenotypes still require elucidation. Further investigations in this field will not only provide insights into bacterial pathogenesis but also shed light on microbial flora-host interactions.

## MATERIALS AND METHODS

### Quorum sensing inhibition assays

The DDR was dissolved in DMSO to obtain a stock concentration of 50 mg μg mL^-1^. Overnight cultures of PAO1-*lasB*-*gfp*, PAO1-*PqsA*-*gfp*, and PAO1-*rhlA*-*gfp* strains were subsequently inoculated in ABTGC medium (0.1% MgCl_2_, 0.1% CaCl_2_, 0.1% FeCl_3_, 10% A10, 0.2% glucose and 0.2% casamino acids) at a dilution ratio of OD_600_ = 0.01. Subsequently, various concentrations of DDR ranging from 5 μg mL^-1^ to 100 μg mL^-1^ or DMSO (0.1%) were added to each well followed by incubation at 37℃ in the Tecan Infinite™200 Pro plate reader (Tecan Group Ltd., Mannedorf, Switzerland) for the measurement of OD_600_ and GFP fluorescence (excitation at 485 nm, emission at 535 nm) with intervals of every15 minutes over a period of 20 hours (hs). The data was calculated as GFP fluorescence/OD_600_.

### Elastase assay

The overnight cultures of PAO1 and the Δ*lasI*Δ*rhlI* mutant negative control were diluted in ABTGC medium to an optical density of 0.01 at OD_600_. DDR (50 μg mL^-1^) and DMSO were added to each well, followed by incubation for 24 hs at 37℃. Subsequently, the culture was centrifuged at 10,000 rpm for 10 minutes, and the supernatants were collected for elastase detection using the EnzChekElastase assay kit (Invitrogen, USA), following the manufacturer’s instructions. The fluorescence signal was measured using a Tecan Infinate 200 Pro plate reader (Tecan Group Ltd., Mannedorf, Switzerland) with excitation at 490 nm and emission at 520 nm.

### Pyocyanin quantification assay

The OD_600_ of overnight cultures of PAO1 and the Δ*lasI*Δ*rhlI* mutant were adjusted to 0.01 in King’s medium supplemented with DDR (50 μg mL^-1^) or DMSO. After incubation at 37℃ for 24 hs, the OD_600_ values were measured and the cultures were centrifuged at 10,000 rpm for 10 minutes. Subsequently, the supernatants were extracted using chloroform (3 mL) and 0.2 M HCl (1.5 mL). The upper aqueous layer containing pyocyanin was transferred into a 96-well microtiter plate for absorbance measurement at 520 nm. The obtained data from OD_520_ measurements were normalized by dividing them with the final OD_600_ values.

### Rhamnolipid quantification assay

The OD_600_ of overnight cultures of PAO1 and Δ*lasI*Δ*rhlI* mutant was standardized to 0.01 in ABTGC medium. Compound DDR (50 μg mL^-1^) or DMSO was added to each group and incubated for 24 hs at 37℃. After determining the OD_600_ of the resulting cultures, supernatants were centrifuged at 10,000 rpm for 10 minutes and extracted twice with diethyl ether. The organic fractions were concentrated to yield white solids, which were then resuspended in deionized water and mixed with a solution containing 0.19% (w/v) orcinol in 50% H_2_SO_4_. Subsequently, the resulting mixture was cooled to room temperature after being incubated at 80℃ for 30 minutes. The absorbance was measured at a wavelength of 421 nm, and the data were normalized using the corresponding OD_600_ values.

### Siderophore assay

For the siderophore assay, bacteria were cultured overnight at 37℃ and then suspended in ABTGC medium with DDR (50 μg mL^-1^) added or not to achieve a final optical density of OD_600_ equal to 0.01 in a 96-well microtiter plate. Subsequently, the plate was placed into a Tecan Infinate 200 Pro plate reader (Tecan Group Ltd., Mannedorf, Switzerland; excitation wavelength: 398 nm, emission wavelength: 460 nm) for fluorescence and OD_600_ measurements. Pyoverdine production was quantified by calculating the ratio of fluorescence to OD_600_ values.

### Intracellular survival and virulence assay

RAW264.7 cells were seeded into 96-well plates with Dulbecco’s modified Eagle’s medium (supplemented with 10% heat-inactivated fetal bovine serum) and incubated for 24 hs. Subsequently, the cells were infected with mid-log phase PAO1 cultured in ABTGC medium with DDR (50 μg mL^-1^) added or not at various multiplicities of infection for 1 h. Following bacterial internalization, the cells were washed twice with pre-warmed phosphate-buffered saline (PBS) to remove extracellular bacteria and then incubated in DMEM medium containing Gm 200 to eradicate extracellular organisms for 1 hour. The GM 200 medium was subsequently replaced with Gm 100 and maintained for an additional 3 hs. Supernatants were collected for lactate dehydrogenase (LDH) detection according to the manufacturer’s instructions (YEASEN, 40209ES76). Cells were rinsed once with PBS, lysed using 0.1% Triton X-100, and plated onto LB agar plates for colony forming unit (CFU) counting.

### *G.mellonella* infection assay

The *G. mellonella* infection assay was performed according to previously published methods (61). Overnight cultures of PAO1 were suspended in ABTGC medium for 6.5 hs, followed by centrifugation at 12,000 rpm and two washes with PBS. Subsequently, the OD_600_ of each group was adjusted to 1 using PBS buffer. *G. mellonella larvae* were randomly assigned to different treatment groups (DMSO, NE, DDR, NE + DDR, Blank), as well as a PBS control group, with each group consisting of 10 larvae. Infection was induced by injecting 10 µL of PAO1 suspension (containing 2×10^4^ CFU mL^-1^) into the left posterior gastropod of each larva. Following injection, the larvae were incubated at 25℃ and monitored at various time points. Larvae were considered dead when they showed no response to external stimuli and displayed dark pigmentation due to melanization.

### Acute lung infection in mice

The infection model was established as previously described (5). Briefly, 7-9-week-old female Balb/c mice were intraperitoneally injected with NE (2 mg kg^-1^), DDR at a dose of 0.67 mg kg^-1^, or a combination of NE (2 mg kg^-1^) and DDR (0.67 mg kg^-1^). Subsequently, the mice were anesthetized with isoflurane and infected intranasally with a dose of 2 × 10^7^ CFUs. After 24 hs of infection, both the non-infected control groups and infected group consisting of six mice each were euthanized. The lungs were then immediately removed and fixed in 4% paraformaldehyde for subsequent sectioning and HE staining. Mouse experiments were conducted in accordance with the guidelines approved by the Animal Subjects Ethics Sub-committee (ASESC) of Hong Kong Polytechnic University (Case No: 23-24/1018-OTHERS-R-NSFC).

### Fluorescence reporter assay

Plasmids containing *gfp* fusions to *cdrA*, *rsmY* or *rsmZ* were transformed into wild-type PAO1. Subsequently, the overnight cultures were diluted to OD600 = 0.01 and transferred into a 96-well microtiter plate with DDR (50 μg mL^-1^) added or not. Finally, the plate was incubated in a Tecan Infinate 200 Pro plate reader (Tecan Group Ltd., Mannedorf, Switzerland; excitation at 488 nm, emission at 535 nm) at 37 °C for 24 hs to monitor fluorescence intensity and OD_600_ values. The obtained data was normalized with respect to OD_600_ values.

### Real-time qPCR

The total RNA from the treatment group exposed to DDR (50 μg mL^-1^) and the blank control cultured in ABTGC medium was extracted using the RNeasy mini kit (Qiagen, Germany), followed by DNase treatment (Qiagen, Germany). Reverse transcription of cDNA was performed using TaKaRa reverse transcription kits. RT-qPCR assays were conducted using Hieff qPCR SYBR green master mix (Yeasen) and qPCR systems from Roche according to the manufacturers’ instructions. The relative expression levels of target genes were determined using the ΔΔCt method after normalization with housekeeping gene *rpsL* values. Data analysis was carried out using LightCycler 96 software from Roche.

### Motility assay

For the swarming motility assay, 2 μL of the overnight bacterial culture was spotted onto the center of Petri dishes containing LB broth agar (0.5%) with corresponding compounds added or not. Swimming and twitching motility assays were performed on LB broth plates containing either 0.3% or 1% agar, respectively. The culture was stab-inoculated into the center or bottom of each respective plate using a toothpick as an inoculation tool. Colony diameter measurements (cm^2^) were taken after incubating all plates at 37 °C for 18 hs.

### Construction of plasmid

DNA cloning and plasmid preparation were performed following standard protocols. PCR primers were designed with restriction sites at their ends for subsequent digestion and ligation into the specific vector (Table S1). For the construction of pET-22b-RetS (residues 41-185), the RetS (residues 41-185) fragment was amplified from genomic DNA using the primer pair 22bRetSf and 22bRetSr. For the construction of pET-28a-N-SUMO-RetS (residues 41-185), the RetS (residues 41-185) fragment was amplified using the primer pair sRetSf and sRetSr. The vector was digested with BamHⅠ and HindⅢ restriction enzymes, followed by purification. The PCR products were then separately digested and inserted into pET-22b or pET-28a-N-SUMO-RetS to generate pET-22b-RetS(residues 41-185) or pET-28a-N-SUMO-RetS (residues 41-185) using a Seamless cloning kit. The resulting plasmids were transformed into *E.coli DH5α*, and their constructs were verified by sequencing using primers that annealed to sites outside the multiple cloning site. For RetS point mutations, PCR was performed using constructed pET-28a-N-SUMO-RetS (residues 41–185) as a template along with corresponding primers. All plasmids underwent verification through DNA sequencing analysis.

**Table S1:**
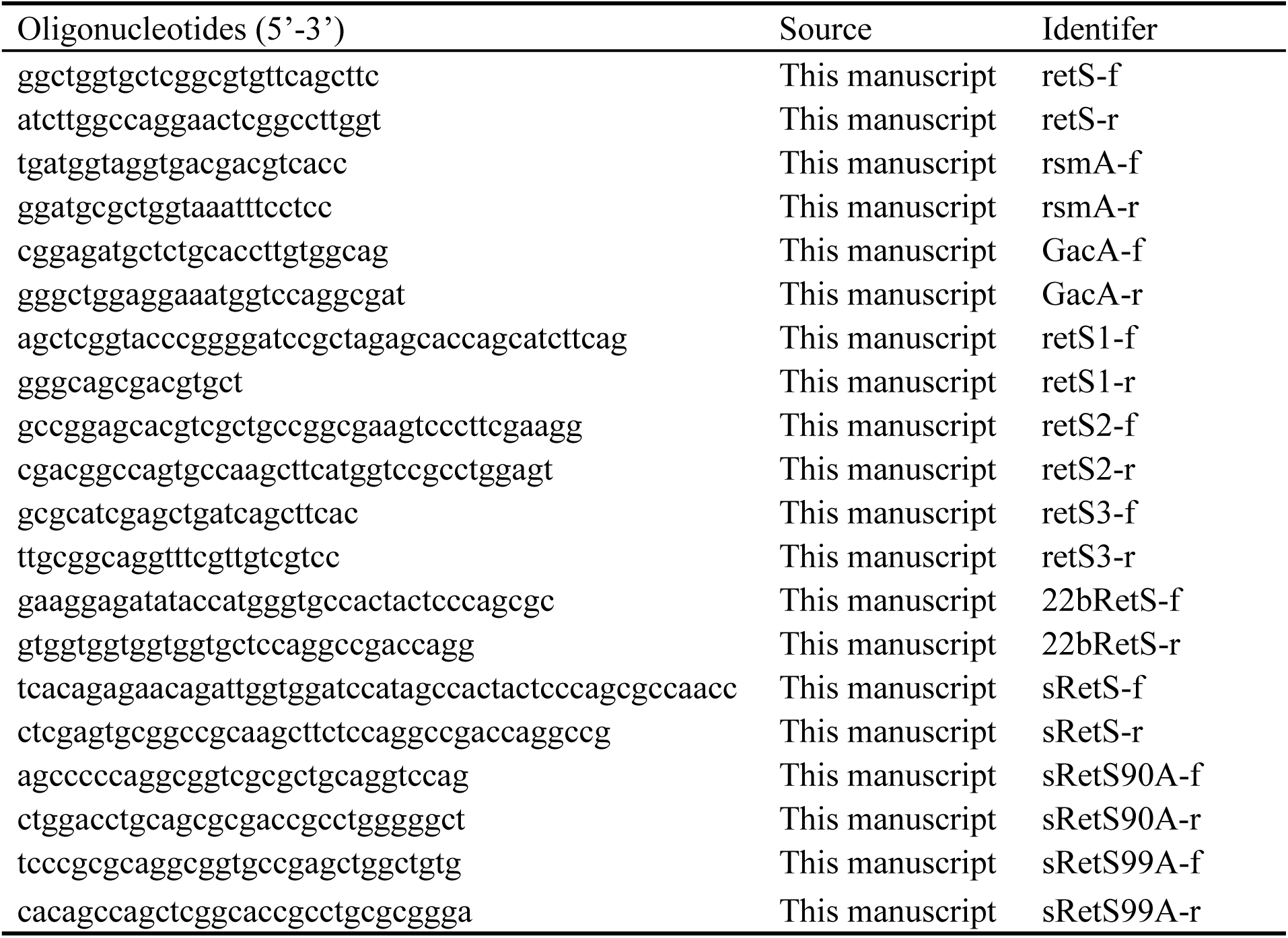

### Protein expression and purification

The plasmid pET-22b-RetS (residues 41-185) or pET-28a-N-SUMO-RetS (residues 41-185) was transformed into *E. coli BL21 (DE3)*, enabling the expression of RetS (residues 41-185) protein containing a 6-His tag. RetS (residues 41-185) was induced with 0.5 mM IPTG overnight at 16 ℃. Cells were harvested and resuspended in lysis buffer composed of 30 mM HEPES pH 6.8 and 150 mM NaCl. Bacteria were lysed using probe sonication for a duration of 40 minutes, followed by removal of cell debris through centrifugation at a speed of 15,000 rpm for 30 minutes. The supernatant was loaded onto a Ni-NTA column (L00250-50, GeneScript). The resin was washed with lysis buffer supplemented to 90 mM imidazole and subsequently eluted with lysis buffer supplemented to 300 mM imidazole. Where necessary, proteins were concentrated via filtration using centrifugation and either a 10 kDa or 30 kDa cut-off.

### Isothermal titration calorimetry (ITC) assays

The isothermal titration calorimetry experiments were conducted at 25 ℃ using a MicroCal PEAQ-ITC instrument (Malvern Panalytical). The protein and ligand solutions were prepared in the same buffer. For the RetS (41–185) - His protein titration experiments with a SUMO tag against NE, the buffer solution comprised of 20 mM HEPES, 150 mM NaCl, pH 6.8, supplemented with 0.1% DMSO. In the RetS (41–185) - His protein titration experiments with a SUMO tag against DDR, the buffer solution also contained 20 mM HEPES, 150 mM NaCl, pH 6.8, however, it included additional components such as 0.1% DMSO, 0.5% Tween-80, and 2% PEG300. A concentration of 67.4 μM RetS (41–185) - His protein or its mutants along with 674 μM NE or DDR was utilized for these assays. Each injection consisted of injecting a volume of 2 μL for a total of twenty injections, the first injection was excluded during data analysis. Data analysis was performed using MicroCal PEAQ-ITC Analysis software (Malvern Panalytical).

### Construction of *P. aeruginosa* mutant strain

The upstream and downstream regions of the *retS* gene were amplified using two pairs of corresponding primers, followed by ligation to the suicide vector PK18 utilizing Gibson Assembly master mix (New England Biolabs [NEB]). Subsequently, the ligated PK18 plasmid (maintained in *E. coli Top10*) was transferred to PAO1 through conjugal mating with assistance from *E. coli* harboring pRK600 vector after sequence verification. Gentamicin was employed for selection of the first homologous recombinants, which were subsequently streaked onto LB agar containing 20% sucrose to select for the second homologous recombinants. Finally, PCR with appropriate primers confirmed the presence of *retS* mutation.

### Crystallization, data collection

The fusion protein RetS41-185 with a N-terminal hexahistidine tag was exogenously expressed by fusing the CDS fragment encoding truncated RetS41-185 to the pET22bN vector. Subsequently, the recombinant plasmid was transformed into *E.coli* [BL21(DE3), CN]. The target protein was purified from the supernatant using Ni-NTA affinity chromatography followed by size exclusion chromatography column. The purity of the protein was confirmed by SDS-PAGE and quantified using NanoDrop One (Thermo Fisher, USA). The high-purity fusion protein was concentrated to 8 mg mL-1 using an ultrafiltration device (Millipore, Germany) and incubated with equimolar NE at a ratio of 1.2:1 for 0.5 h. Subsequently, 1 μL of the post-incubated protein was mixed with an equal volume of reservoir solution and then equilibrated over 100 μL of reservoir solution containing 0.1 M Bis-Tris Propane pH 8.5, 0.2 M NaI, and 20% polyethylene glycol (PEG) 3350 in a sealed cell at a temperature of 16℃ using the vapor diffusion-sitting-drop method. To collect diffraction data, the crystal was picked and flash-cooled in liquid nitrogen along with the reservoir solution containing 20% glycerol as a cryoprotectant before being shipped to Shanghai Synchrotron Radiation Facility (SSRF). Finally, single-crystal diffraction was performed by exposing it to synchrotron radiation beam with a wavelength of 0.97946 Å while maintaining it at a temperature of 100 K in the stream of liquid nitrogen using small angle oscillation method at BL18U1 in SSRF. Meanwhile, synchronized recording of diffraction frames was carried out using a PILATUS3 detector with each frame captured every degree.

### Data processing, phasing, model building and refinement

Firstly, all raw diffraction frames were sequentially indexed and integrated using XDS with the assistance of XDSGUI. Subsequently, the resulting integral file named XDS_ASCII.HKL was subjected to further processing in Aimless package within CCP4 Suite, leading to a merged scaled file with an mtz suffix. Following this, molecular replacement was performed utilizing PHASER in Phenix by employing the crystal structure of apo PaRetSp (PDB accession code: 3JYB) as a search template. Model building was then carried out using Autobuild in Phenix. Finally, COOT and Phenix were employed for refining and completing the initial model respectively, while Molprobility in Phenix served as a validation tool. Detailed information regarding data collection and processing can be found in Table S2. The final atomic coordinates and structure factor files have been deposited in the Protein Data Bank under accession code 9JPE.

**Table S2:**
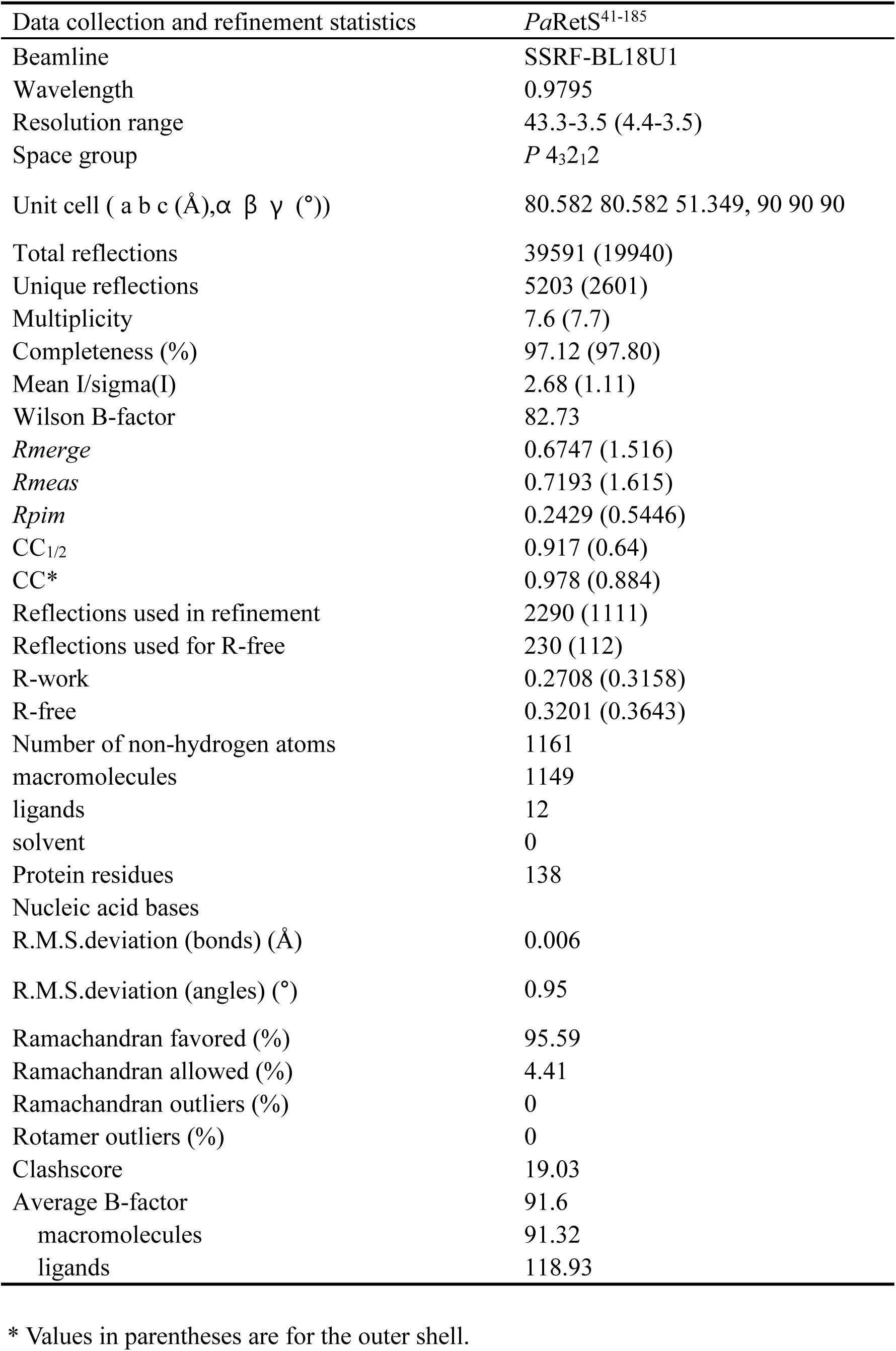

### Statistical analysis

Data were presented as the mean ± S.D. P values < 0.05 were considered significant. Statistical analysis and quantitative data plotted were performed by Graph pad Prism software with one-way ANOVA analysis. All experiments were conducted at least three replicates.

## Data, Materials, and Software Availability

All study data are included in the article and/or supporting information.

## ACKNOWLEDGMENTS

This work was supported by the National Key Research and Development Program of China (2022YFC2304700); the National Natural Science Foundation of China (32270196 and 32200053); Shenzhen Science and Technology Program (KQTD20200909113758004); HaiYa Young Scientist Foundation of Shenzhen University General Hospital (2024-HY013); and Guangdong Pearl River Talent Plan (2019QN01Y163).

## Author contributions

Y.Z., performed most experiments wrote the manuscript; A.R., conduct ITC assay; C.W., performed molecular spectral analysis. T.Z., contributed to project discussion, J.D., donate retS mutant. Y.H and R.C assisted with preparation of reagents. W.H., W.Z and Y.F provided conceptual advice. H.L reviewed and edited this manuscript. L.Y acquires the funding, supervised the conduction of project, review and edited this manuscript.

